# Topotecan Synergizes with SP141 in the Management of Uveal Melanoma

**DOI:** 10.64898/2026.09.23.753860

**Authors:** Anju Thomas, Utkarsh R. Addi, Audrianna Wu, Kaleigh Kozak, Ross F. Collery, Amit Joshi, Aparna Ramasubramanian, Shyam S. Chaurasia

## Abstract

**Purpose:** Uveal melanoma (UM) is a rare but aggressive intraocular malignancy originating in the uvea (choroid, iris, and ciliary body). Despite primary treatment, 50% of UM patients develop metastases to the liver, lungs, bones, and skin. Topotecan, a topoisomerase I inhibitor, and SP141, an MDM2 inhibitor, have been studied in cancers, including retinoblastoma. This study explores a combined approach with these drugs in the 92-1 UM cell line.

**Methods:** Primary choroidal melanoma-derived 92-1 cells were treated with topotecan and SP141 individually to determine IC_50_ values. Cell viability and proliferation were assessed using the MTT assay and the xCELLigence real-time monitoring system. The synergistic effects were analyzed quantitatively using the Loewe Additivity and Highest Single-Agent (HSA) models. Flow cytometry was employed to analyze cell cycle distribution, while cell death and apoptosis were monitored via high-throughput, real-time quantitative analysis using the IncuCyte live-cell kinetic imaging system.

**Results:** Both Topotecan and SP141 induced significant cell death in 92-1 cells. xCELLigence showed dose- and time-dependent inhibition of proliferation. Combining sub-IC_50_ concentrations of topotecan and SP141 exhibited significant synergistic effects, markedly reducing growth. Cell cycle analysis revealed that topotecan causes a dose-dependent biphasic arrest in G2 and S phases, while SP141 causes G2/M arrest, leading to cell death. Cytotoxicity assays corroborated increased cell death following combination therapy, and the Caspase 3/7 assay indicated that apoptosis is the mechanism underlying cell death.

**Conclusions:** The multi-targeted approach combining topotecan and SP141 drugs demonstrated synergistic therapeutic effects in 92-1 cells, suggesting a potential treatment strategy for managing UM.

## Introduction

Uveal melanoma (UM) is the most common primary intraocular malignancy in adults.^1^ It accounts for 5-10% of all melanoma cases, with an incidence of roughly 4.3 per million people per year.^2^ UM arises from melanocytes in the uveal tract (most commonly the choroid), but its biology and progression differ considerably from those of skin melanoma. The 5-year survival rate for skin melanoma, based on SEER (Surveillance, Epidemiology, and End Results) data, is 94.7%, compared with 82.8% for uveal melanoma.^3,4^ Importantly, the mortality rate from cutaneous melanoma has significantly improved from 1975 to 2019.^5^ The current 5-year survival statistic is largely attributable to the introduction of multiple new, highly efficacious treatments for cutaneous melanoma in 2013. Contrastingly, UM treatment options and 5-year survival have remained largely unchanged since 1973.^5^ Current UM patients are, therefore, left with limited treatment options known to present a significant risk of complications.

UM is primarily treated with plaque brachytherapy, first described by Stallard in the 1960s.^6^ Advanced UM is treated mainly with enucleation, and secondary enucleation may be performed for tumor recurrence, non-response, or neovascular glaucoma.^7^ A large prospective clinical trial – the Collaborative Ocular Melanoma Study (COMS) showed that large- and medium-sized choroidal melanomas treated with enucleation or brachytherapy had similar 5-year survival rates.^8^ Though plaque brachytherapy preserves the eye globe, it has demonstrated a 10% treatment failure rate and is associated with a significant decline in visual acuity. In the COMS study, substantial visual impairment was defined as a loss of 6 or more lines of visual acuity from the pretreatment level (49% of eyes) or a visual acuity of 20/200 or worse (43% of eyes) 3 years after iodine-125 brachytherapy.^9^

Regardless of the primary treatment method, about half of UM patients develop metastases to other parts of the body.^10^ The most common sites of metastasis are the liver (95%), lungs (24%), bone (16%), and skin (11%).^11^ There is no current effective treatment for metastatic disease, resulting in a high mortality rate among patients with UM.^12^ Therefore, there is an unmet need for new treatment strategies for UM that provide effective local control, minimize visual acuity loss, and reduce the risk of metastatic spread.

Topotecan hydrochloride is a topoisomerase I inhibitor that prevents the re-ligation of single-strand DNA breaks during replication, induces DNA damage and replication stress, and thus activates apoptosis in rapidly dividing tumor cells.^13–15^ It serves as a chemotherapy drug in several cancers, including its local use in retinoblastoma, with minimal retinal toxicity.^16–18^ Murine Double Minute 2 (MDM2) inhibitors, on the other hand, are a class of drugs that disrupt the interaction between the E3 ubiquitin ligase and the tumor suppressor gene, p53. Under physiological conditions, MDM2 negatively regulates p53 by promoting its ubiquitination and proteasomal degradation, thereby limiting p53’s ability to induce cell-cycle arrest. Inhibition of MDM2 stabilizes wild-type p53, inducing its tumor-suppressor function.^19,20^ In UM, *TP53* mutations are rare, so targeting MDM2 to restore p53 tumor-suppressor function can be an effective strategy to induce cell-cycle arrest, particularly in cancers that retain functional p53. Previously, MDM2 inhibitors have been studied in preclinical models in UM. For example, nutilin-3 and CGM097 have been shown to activate p53 signaling and demonstrate antitumor effects primarily as monotherapies.^21,22^ Other MDM2 inhibitors, such as HDM201 and alrizomadin (APG-115), demonstrated synergism with PKC and BCL2 inhibitors, supporting the feasibility of combination strategies for UM.^23–25^ Topotecan has been studied in combination with the MDM2 inhibitor nutilin-3 in UM cell lines^26^ to inhibit growth and promote apoptosis, suggesting that combining DNA-damaging agents with MDM2 inhibitors could be therapeutically promising. Nevertheless, the high dosage requirement, dose-limiting toxicity, and limited bioavailability of nutlin-3 have rendered it suboptimal for clinical application.^27^ The present study explores the therapeutic potential of SP141, a recently developed MDM2 inhibitor that directly and specifically binds MDM2 and inhibits its activity, thereby reactivating p53 tumor-suppressor function, a mechanism that remains largely unexplored in UM. Additionally, the study will evaluate the multi-target and synergistic effects of the chemotherapy drug topotecan in combination with SP141 in 92-1 cells.

## Materials and Methods

### Cell culture and Reagents

92-1 cells were selected for these studies because previous reports have indicated their reliability in assessing the genetic, phenotypic, and metastatic traits of primary human UM^28–30^. In brief, the 92-1 cell line was established from a primary choroidal melanoma in a 76-year-old female patient at the Leiden University Medical Center.^30^ It harbors the GNAQ mutation, which activates the Gαq pathway, and the *EIF1AX* mutation.^28,31^ Since 85-90% of UM tumors originate from the choroid,^11^ this established cell line is frequently used as an in vitro model for UM.^21–23,26^

92-1 cells were procured (Cat# 13012458, Millipore Sigma, Burlington, MA) and cultured in RPMI 1640 media (Thermo Fisher Scientific, Waltham, MA) supplemented with 10% FBS (MIDSCI, Omaha, NE) and 1X Antibiotic Antimycotic (Thermo Fisher Scientific) in a humidified atmosphere at 37°C with 5% CO_2_. Topotecan was obtained from Hospira, Inc. (Cat# NDC-0409-0302-01, Lake Forest, IL). 1 mg/ml stock solutions were prepared in sterile saline, aliquoted, and stored at -20°C for long-term storage. SP141 was supplied by 3T Ophthalmics (Irvine, CA), and 1 mg/ml stock solutions were prepared in DMSO, aliquoted, and stored at -20°C. The human-derived immortalized retinal pigment epithelium (RPE) cell line (ARPE-19, ATCC® CRL2302™, Manassas, VA, USA) was used to assess the toxicity, tolerability, and potential side effects of topotecan and SP141 (both in mono- and combination therapy studies).^32,33^ ARPE-19 cell line authentication was achieved through Short Tandem Repeat (STR) profiling and multiplex PCR (IDEXX BioAnalytics, Columbia, MO). ARPE-19 cells were cultured in DMEM: F12 (ATCC 30-2006) supplemented with 10% FBS (ATCC 30-2020) and penicillin/streptomycin antibiotics, as described previously.^34^

### Real-Time Cellular Analysis

92-1 cells were continuously monitored for cell behavior, including proliferation, viability, morphology, and adhesion, using the automated xCELLigence real-time monitoring system (Agilent Technologies Inc., Santa Clara, CA). Cells were seeded at a density of 10,000 cells per well in an E-plate with gold electrodes (Cat# 5469830001, Agilent, Santa Clara, CA), and treatments were performed after 24 h. Topotecan treatments were performed at concentrations of 10 nM, 25 nM, 50 nM, 100 nM, 200 nM, 500 nM, 1000 nM, 1500 nM, 2000 nM, and 5000 nM, while SP141 was tested at concentrations of 250 nM, 500 nM,1000 nM, 1500 nM, 2500 nM, 5000 nM, 10,000 nM and 20,000 nM to evaluate the optimal dose.

### Cell Viability Assay

92-1 cells were seeded in 96-well plates (2500 cells/well) and treated with topotecan and SP141 for 24 h, 48 h, 72 h, and 96 h. Cell viability was assessed at each time point using the MTT assay (Thermo Fisher Scientific, Carlsbad, CA). Media (100 µL/well) containing 10 µL MTT (3-(4,5-dimethylthiazol-2-yl)-2,5-diphenyltetrazolium bromide) was aspirated from each well. Plates were incubated for 3 h at 37°C, followed by aspiration of 80 µL of MTT solution, addition of 50 µL of DMSO per well to dissolve the formazan crystals, and a 10 min incubation at 37°C. Absorbance was measured at 540 nm (BioTek Instruments, Winooski, VT). IC_50_ values for topotecan and SP141 were determined at 72 h. Results were analyzed, and IC_50_ values were calculated using the variable-slope method in GraphPad Prism Version 10.6.1 software (GraphPad Software, Boston, MA).

### Cell Cycle Analysis

92-1 cells were seeded in 100-mm dishes (0.5 x 10^6^ cells/dish) and harvested after 72 h of treatment with topotecan and SP141, fixed with chilled absolute ethanol by vigorous vortexing, and stored overnight at -20°C. Cells were washed and then incubated with PI staining buffer (BD Biosciences, Franklin Lakes, NJ) and 100 μg/ml RNase A (Sigma-Aldrich, St. Louis, MO) at room temperature for 1 h in the dark. Cells were filtered through a 30 μm nylon-mesh filter and suspended in PBS. Flow cytometry was performed on an LSRFortessa X20 flow cytometer (BD Biosciences, San Jose, CA). A total of 100,000 events were recorded using appropriate stop gates set in the PI-fluorescence area, compared with a width projection to exclude cell aggregates. Results were analyzed using FlowJo v10.7.1 (BD Biosciences, Ashland, OR).

### Drug Synergism

The multi-targeted therapeutic effects of Topotecan and SP141 were evaluated using MTT cell viability assay. Data were imported into Combenefit software (v2.02) and evaluated using two approaches. The Highest Single Agent (HSA) model compares combined inhibition with the most effective individual drug effect. It calculates synergistic effects by mapping data, fitting dose-response curves, and computing excess inhibition relative to the maximum single-agent inhibition, with heatmap visualizations.^35^ The Loewe additivity model assesses drug interactions by comparing the effect of a combination with that of a single drug when the drugs act on the same pathway. It defines synergism as a situation in which a combination requires lower doses than would be expected under additivity.^36^ A positive score indicates synergism, a score near zero suggests additivity, and a negative score depicts antagonism. Heatmaps were generated to visualize interactions across the tested combinations at several concentrations in this study.

### Real-time Cytotoxicity and Cell Death Assay

92-1 uveal melanoma cells were seeded at 2,500 cells/well in 96-well plates and allowed to adhere for 72 h. Cells were then treated with topotecan (25 nM), SP141 (150 or 600 nM), and their combinations (25 nM topotecan + 150 nM SP141 and 25 nM topotecan + 600 nM SP141). To detect cell death and apoptosis, wells were supplemented with IncuCyte® Cytotox Red Reagent (250 nM; Cat. #4632) and IncuCyte® Caspase-3/7 Green Reagent (5 µM; Cat. #4440) as per the respective assay. Plates were continuously monitored using the IncuCyte SX3 Live-Cell Analysis System (Sartorius Corporation, Bohemia, NY), a turnkey high-performance incubator-based platform designed for automated, kinetic imaging of living cell populations. It provides HD phase-contrast and two-channel fluorescence as readouts to capture cellular movement and molecular changes, enabling analysis of cell morphology and health. Images were acquired over time, and cytotoxicity/apoptosis data were quantified using the IncuCyte Basic Analyzer by calculating fluorescent object counts normalized to cell confluence. Fluorescent objects corresponding to Caspase-3/7-positive cells and Cytotox positive cells were identified using a predefined fluorescence segmentation mask. The segmentation parameters were established using representative control and treated wells and verified to ensure appropriate detection of fluorescent objects while minimizing background signal and cellular debris. The same analysis definition was subsequently applied to all treatment groups and time points.

### Topotecan and SP141 Tolerability

ARPE-19 cells were seeded into 96-well plates at a density of 20,000 cells per well in 200 µL of growth medium. Plates were incubated at 37°C for 24 h to reach ∼90% confluence. To begin drug treatment, topotecan (10-5000 nM) and SP-141 (100-20000 nM) were diluted from stock solutions in the medium. 200 µL medium was removed from each well, and 200 µL of each drug treatment was administered in triplicate. Controls for topotecan and SP141 (pure medium and 250 nM DMSO in medium, respectively) were also administered in triplicate. Treated cells were incubated at 37°C for 72 h. Cell viability was assessed using an MTT assay as described above, in addition to an LDH assay (CyQUANT™ LDH Cytotoxicity Assay Kit, Thermo Fisher Scientific). ARPE-19 cells were cultured with topotecan and SP141, and 110 µL of medium was removed from each well at 72 h, then stored for the LDH assay in accordance with the manufacturer’s instructions. Briefly, 50 µL of sample medium from the 72 h drug treatment plate was added to each well in a 96-well plate, and 50 µL of 1X LDH Positive Control was aliquoted into triplicate wells. 50 µL of the reaction mixture was added to each well, and the samples were mixed thoroughly. The plate was covered and incubated at room temperature for 30 min. 50 µL of the stop solution was added to each sample well, and the wells were gently tapped to mix. The absorbance was measured at 490 nm and 680 nm (BioTek Instruments, Winooski, VT).

### Statistical analysis

All experiments were performed in triplicate and independently repeated twice, unless otherwise specified. Differences among treatment groups were evaluated using one- or two-way analysis of variance, depending on whether a single or two independent variables affected a single dependent variable. Subsequent post hoc analyses were conducted using GraphPad Prism Ver 11.0.2 (GraphPad Software, Boston, MA). Results with a p-value less than 0.05 were considered statistically significant. Data are expressed as mean ± SD.

## Results

### Topotecan and SP141 induce cytotoxic morphological changes in 92-1 UM cells

Topotecan treatment at concentrations of 0-100 nM altered cell morphology and density in 92-1 cells after 72 hours (Fig. 1A-D). Increasing concentrations of topotecan led to changes in shape, density, and morphology compared with the vehicle-treated control group. Additionally, the cells exhibited shrinkage and rounding, detachment from the surface, and cytoplasmic condensation, suggesting cytotoxicity associated with apoptosis and cell-cycle arrest. Similarly, SP141 induced significant dose-dependent (0-1000 nM) changes in cell shape, size, and structure (Fig. 1E-H), with a lower dose (100 nM) showing initial signs of stress, including rounding and decreased confluence (Fig. 1F). Treatment with 600 nM and 1000 nM resulted in a substantial reduction in cell density, with significant rounding, swelling, cell detachment, and accumulation of cell debris, suggesting cytotoxicity (Fig. 1G-H).

**Figure 1.**
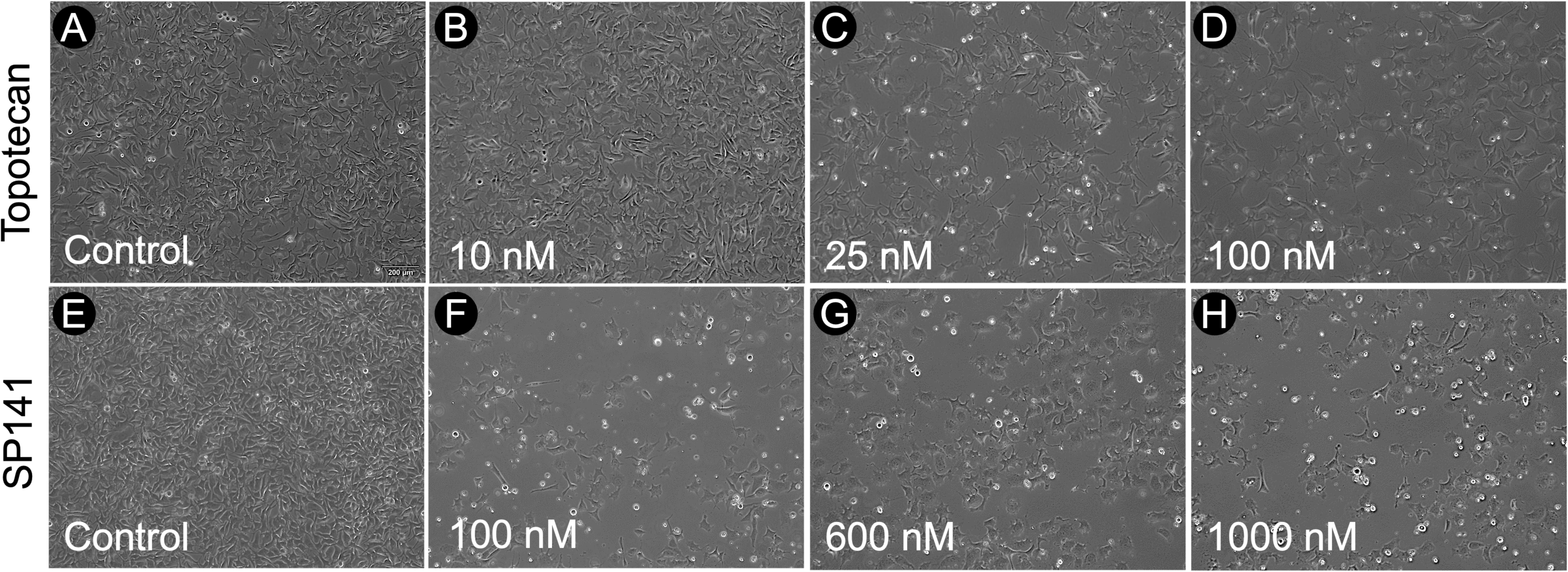
Topotecan and SP141 induce morphological changes in 92-1 UM cells. 92-1 cells seeded at a density of 1 × 10^3^ cells showed altered cell morphology and density after 72 hours of treatment with topotecan at 0-100 nM (Fig. 1A-D) or SP141 at 0-1000 nM (Fig. 1E-H). Higher doses of topotecan (25 and 100 nM) caused cells to change shape, become denser, and exhibit altered morphology, with features such as shrinking, rounding, detachment, and cytoplasmic condensation, suggesting cytotoxicity, apoptosis, and cell-cycle arrest. In the SP141 group, a lower dose (100 nM) led to stress signs like rounding and decreased confluence (Fig. 1F). At higher SP141 doses (600 and 1000 nM), cell density was greatly reduced, with rounding, swelling, detachment, and debris formation, indicating cytotoxic effects (Fig. 1G-H). n=6 in each group and scale bar = 200 μm.

### Topotecan and SP141 inhibit 92-1 UM cell growth

To evaluate the cellular adhesion and proliferation of 92-1 UM cells with topotecan and SP141, increasing concentrations of topotecan (10-5000 nM) and SP141 (250-20,000 nM) were analyzed using a real-time, label-free measurement of electrical impedance across specialized E-Plates integrated with gold microelectrodes using the xCELLigence system. The normalized Cell Index (CI) was monitored over time to compare treatment groups, providing a unitless measure that correlates with cell number, size, and adhesion strength, indicating cellular growth at about 36 hours post-seeding. CI values in controls increase steadily, consistent with exponential cell proliferation, cell spreading, and strong attachment to the substrate (Fig 2A and 2B). Administration of topotecan (Fig 2A) consistently decreases the CI, indicating an inhibition of proliferation, concentration-dependent cell detachment, and possible death in 92-1 cells. In brief, treatment with 10 nM and 25 nM topotecan induced a slight decrease in the CI. However, 50 nM and 100 nM show a cytostatic response 36 h post-treatment, suggestive of slowed or stalled cellular proliferation. At 200-5000 nM, a significant arrest in CI progression was observed at 36 h post-treatment, indicating full suppression of proliferative activity. Topotecan concentrations ≥1000 nM had CI values below baseline, indicative of cytotoxicity.

**Fig. 2.**
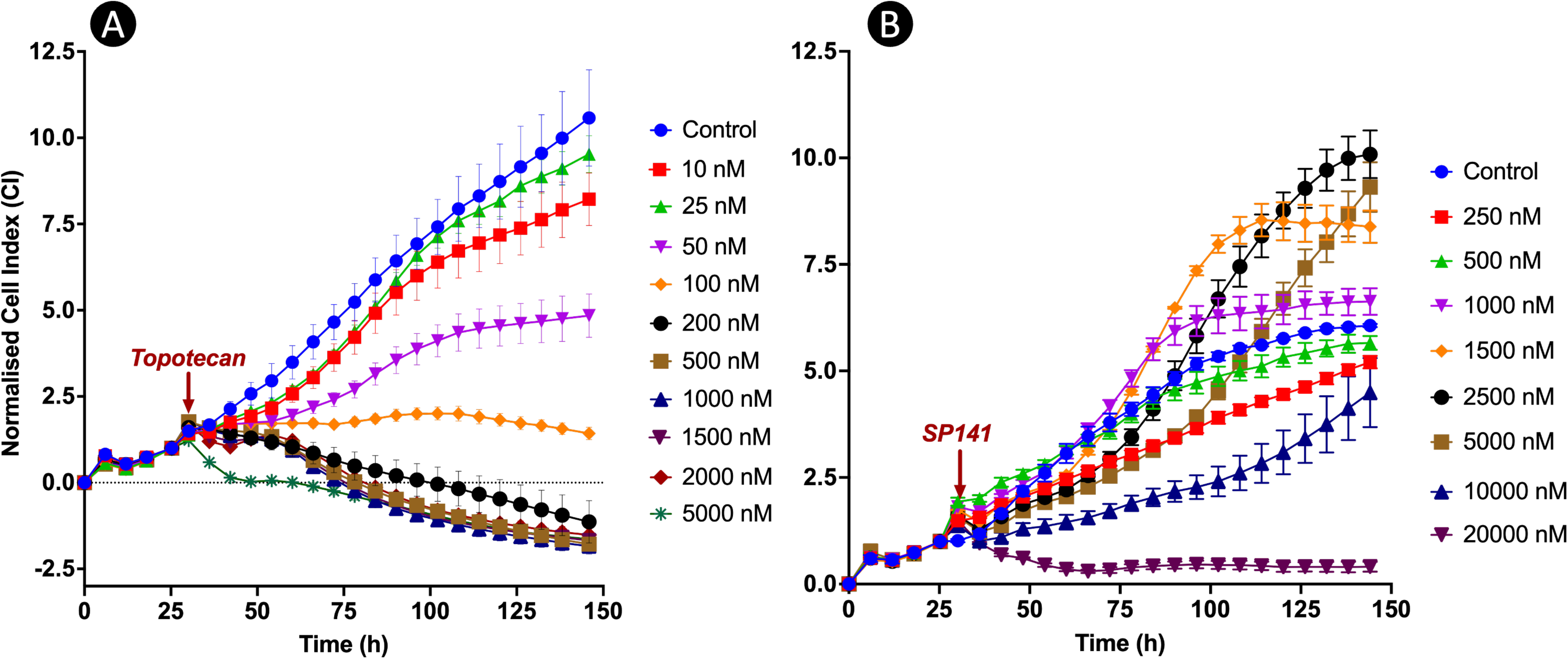
Topotecan and SP141 inhibit the proliferation of 92-1 uveal melanoma cells. Real-time monitoring of 92-1 UM cell adhesion and proliferation after treatment with Topotecan (A) and SP141 (B), using the xCELLigence system. Results are shown as mean ± standard deviation (SD). Each data point was collected in triplicate and analyzed using xCELLigence’s software (Agilent Technologies Inc., Santa Clara, CA). The 92-1 UM cells were monitored for 145 h post-seeding, and the arrows indicate the time point at which the drugs (Topotecan or SP141) were added.

SP141 was assessed similarly and applied 36 h post-seeding (Fig. 2B). CI values in the control group continued to increase through 150 h. SP141 was administered at 36 h. 250 and 500 nM SP141 caused a slight decrease in CI, whereas concentrations of 1500 to 5000 nM resulted in a significant increase in CI after 72 h of treatment. Because CI positively correlates with contact area, the observed increase in CI may reflect cytotoxicity-induced cell swelling, as shown in Figure 1, which increases impedance. However, higher concentrations of 10,000 nM and 20,000 nM caused significant inhibition of proliferation. At these doses, CI values plateaued or declined post-treatment and remained suppressed, indicating anti-proliferative effects and cytotoxicity of SP141 in 92-1 cells.

### Topotecan and SP141 decreased the cell viability of 92-1 UM cells in a dose-dependent manner

To assess the effects of topotecan and SP141 on cell viability and cytotoxicity, 92-1 UM cells were treated with various concentrations of these drugs for 72 h. Cell viability was measured using the MTT assay, and dose-response curves were plotted in GraphPad Prism using a nonlinear regression model with a variable slope and a four-parameter logistic curve to determine IC₅₀ values (Fig. 3A-D).

**Figure 3.**
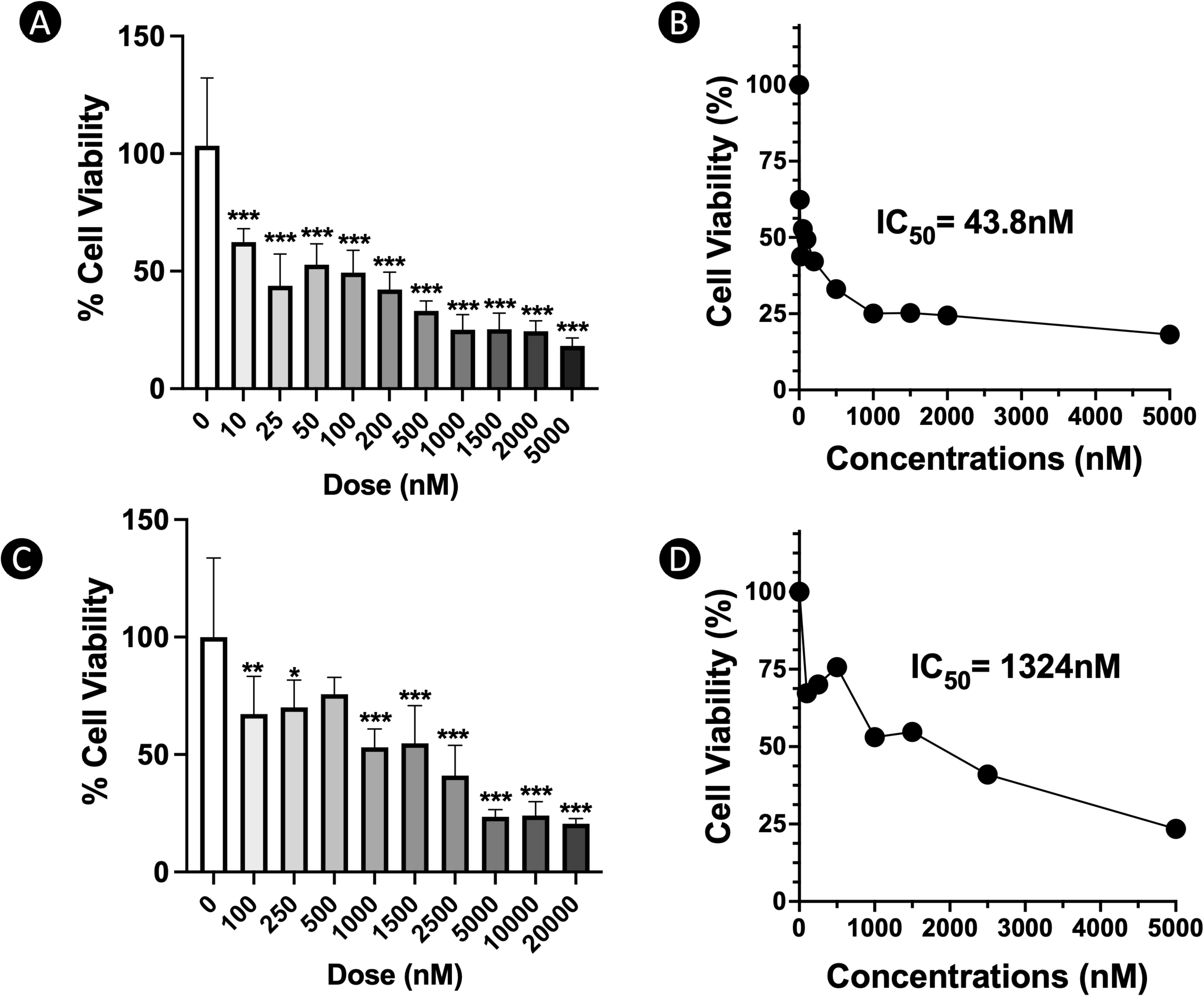
Topotecan and SP141 reduced cell viability in 92-1 UM cells. Bar graph showing percentage cell viability of 92-1 cells measured by MTT assay after treatment with various concentrations of Topotecan (A) and SP141 (C) for 24, 48, 72, and 96 hours. The estimated IC_50_ values for Topotecan and SP141 were 43.8 nM (B) and 1324 nM (D), respectively. Data are represented as mean ± standard deviation (SD). n = 6 in each group. *, p ≤ 0.05; **, p ≤ 0.01; and ***, p ≤ 0.001 compared with the control group (0 nM).

Topotecan treatment resulted in a concentration-dependent and significant reduction in cell viability at 24, 48, 72, and 96 h (Supplementary Fig. 1A). At 24 h, most concentrations of topotecan had minimal impact on the viability of 92-1 cells (Supplementary Fig. 1A). By 48 h, exposure to 25 nM or higher led to a significant decrease in viability, with a 50% reduction at doses above 500 nM. At 72 h, a significant cytotoxic effect was observed as early as 10 nM (Fig. 3A), and viability continued to decrease at higher concentrations, remaining above 50% at ≥ 25 nM. The IC_50_ for topotecan at 72 h was 43.8 nM (Fig. 3B).

Similarly, SP141 treatment significantly decreased cell viability in a concentration-dependent manner at 24, 48, 72, and 96 h. At 24 h, higher doses showed a significant reduction in cell viability (Supplementary Fig. 1B), but by 48 h, concentrations ≥ 300 nM began to significantly reduce cell viability, with a 50% or greater reduction was observed at concentrations > 2.5 µM (Supplementary Fig. 1B). The cytotoxic effect became more evident by 72 h, with a 50% decrease in viability observed at 1000 nM and a significant decline from 5000 nM and above concentrations (Fig. 3C). At 96 h, SP141 continued to decrease cell viability in a dose-dependent manner. The IC_50_ for SP141 at 72 h was 1324 nM (Fig. 3D).

### Topotecan and SP141 arrest 92-1 cells at different cell cycle phases

To understand how topotecan and SP141 influence the cell cycle mechanisms, flow cytometry was used to analyze PI staining and cell cycle distribution after 72 h of treatment. Treatments included topotecan at 25 nM, 45 nM, and 100 nM, and SP141 at 100 nM, 600 nM, and 1000 nM, selected based on previous results (Supplementary Fig. 1) and IC_50_ values (Fig. 3A-D), with each compound tested separately in 92-1 cells.

A normal proliferative profile of 92-1 cells showed 48-55% in G1, 29-31% in S, and 16-20% in G2/M, with no appreciable sub-G1 population in the vehicle-treated control group (Figs. 4A and 5A). The 25 nM topotecan treatment significantly increased the proportion of cells in G2/M phase (46%), suggesting G2/M arrest (Figure 4B, 4F). In the 45 nM topotecan-treated group, the cells showed a shift, with 62% of the population arrested in S phase, an increase in the sub-G1 population (11%), and a notable decrease in both G1 phase (10%) and G2/M phase (16%) (Figure 4C, 4F). Similar observations were made at 100 nM topotecan, where cells in S phase accounted for 61% of the population and 10% were in the sub-G1 fraction (Figure 4D, 4F). The results indicate that topotecan, a known topoisomerase I inhibitor that disrupts replication fork progression, exhibited a biphasic pattern of cell-cycle arrest (G2/M and S phases) in 92-1 cells at increasing concentrations.

**Figure 4.**
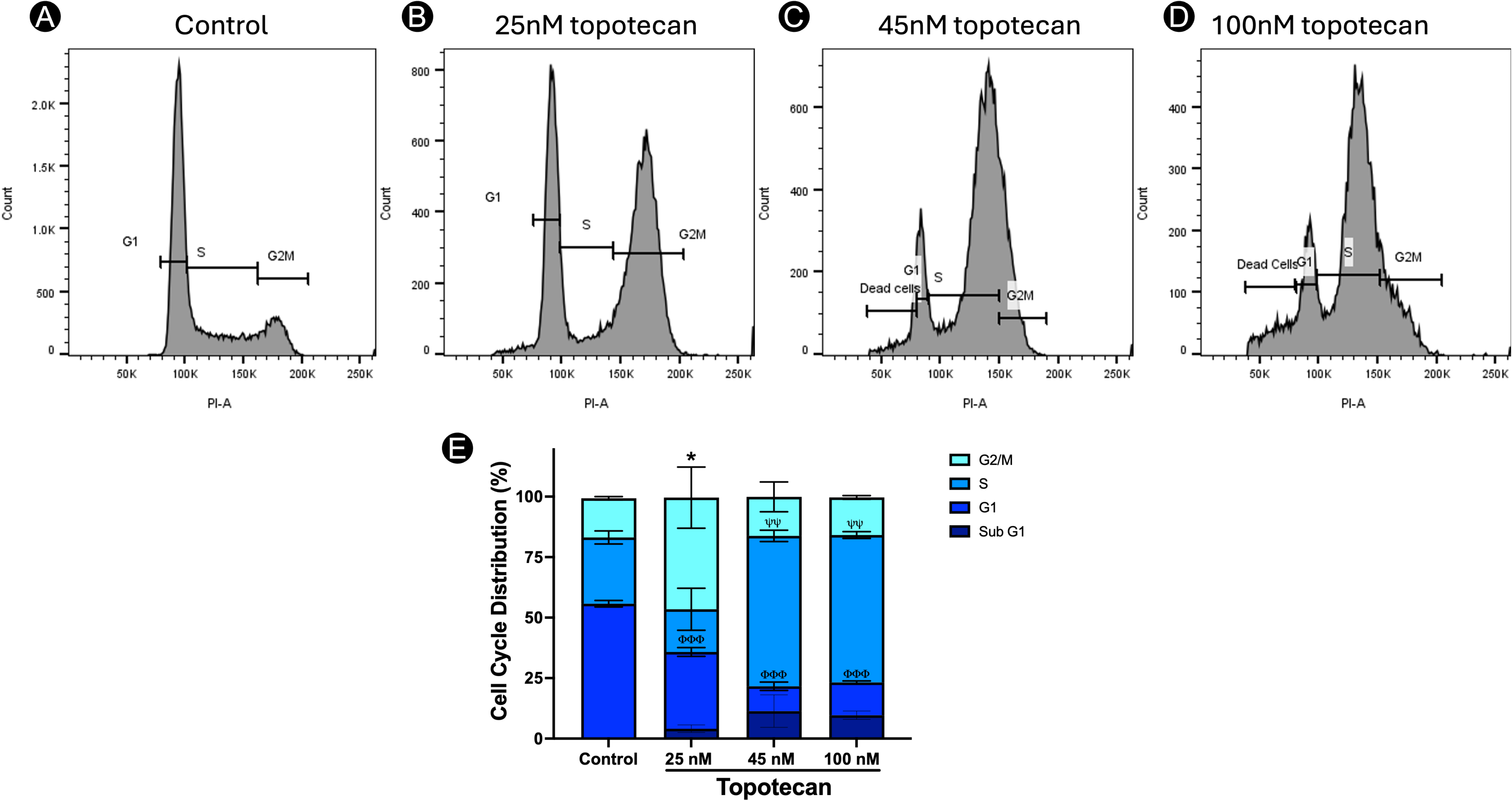
Topotecan arrests 92-1 UM cells in the G2/M and S phases of the cell cycle. Cell cycle analysis histograms of 92-1 UM cells after 72 h of Topotecan treatment (A-D). The cell cycle distribution percentages were calculated and presented as a bar graph showing cell cycle arrest across different phases (E). Data are presented as mean ± standard deviation (SD). *, p ≤ 0.05 compared with the control group in the G2/M phase. ^ψψ^, p ≤ 0.01, ^ψψψ^, p ≤ 0.001 compared with the control group in the S phase. ^ϕϕϕ^p ≤ 0.001 compared with the control group in the G1 phase.

**Figure 5.**
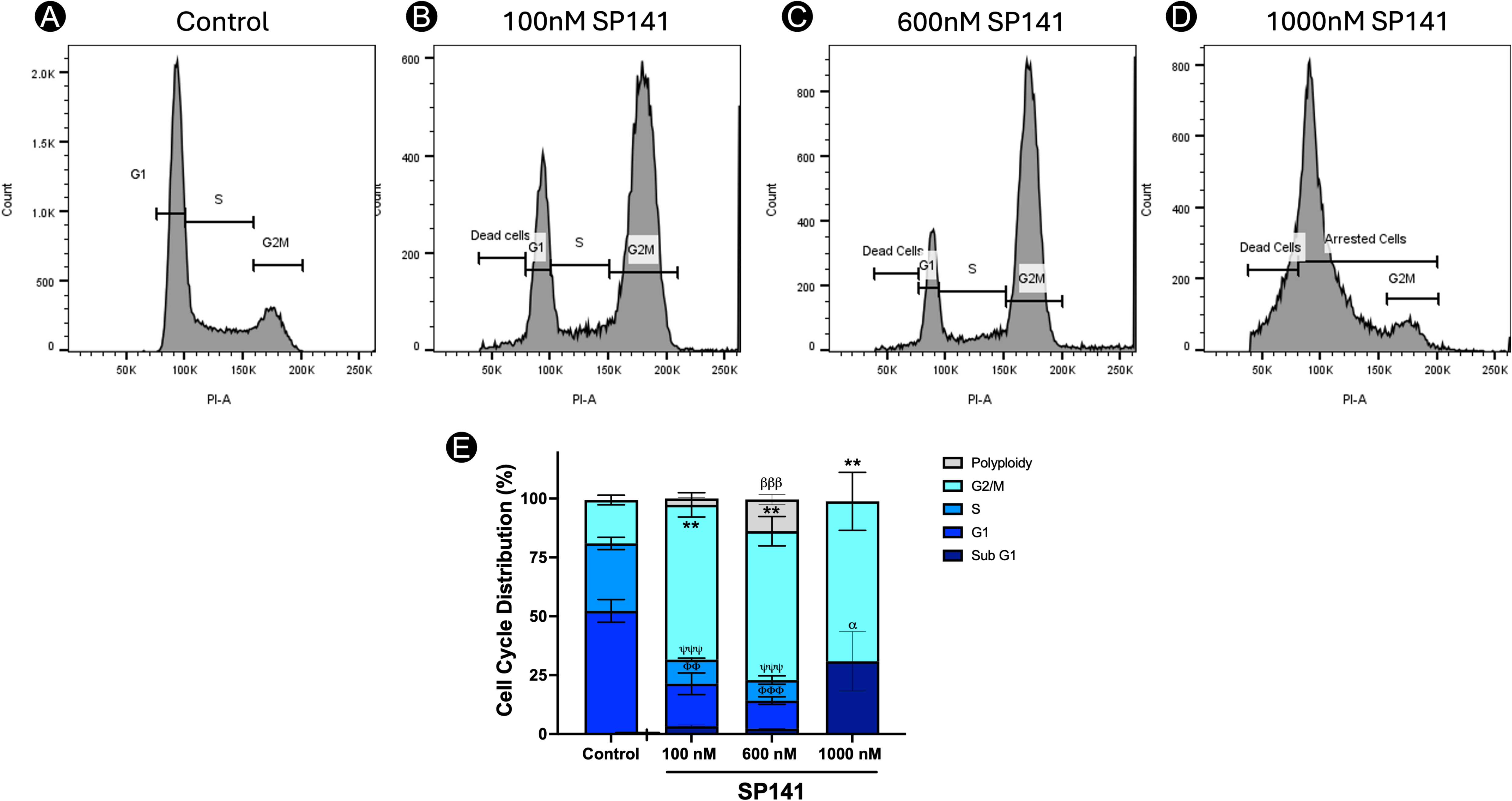
SP141 arrests 92-1 UM cells in the Sub-G1 and G2/M phases of the cell cycle. Cell cycle analysis histograms of 92-1 UM cells after 72 h of SP141 treatment (A-D). The cell cycle distribution percentages were calculated and presented as a bar graph showing cell cycle arrest across different phases (E). Data are presented as mean ± standard deviation (SD). ^βββ^, p ≤ 0.001 compared with the control group for polyploidy. **, p ≤ 0.01 compared with the control group in G2/M phase. ^ψψψ^, p ≤ 0.001 compared with the control group in S phase. ^ϕϕ^, p ≤ 0.01; ^ϕϕϕ^, p ≤ 0.001 compared to the control group in G1 phase. ^α^, p ≤ 0.05 compared with the control group in the sub G1 phase.

On the other hand, SP141 consistently arrested the cell cycle in the G2/M phase. 100 nM SP141 treatment resulted in a significant increase in G2/M phase (66%) and a corresponding decrease in G1 (18%) and S (10%) phases (Figure 5B, 5F). At 600 nM, G2/M arrest was maintained (63%), and polyploid cells were observed (14%), along with a decrease in G1 and S phases (Figure 5C, 5F). At 1000 nM, 31% of the population underwent cell death or entered the Sub-G1 phase, with a significant arrest at G2/M (68%) (Figure 5D, 5F). SP141 induced a dose-dependent cell cycle arrest of 92-1 cells in the G2/M phase, with accumulation of polyploid cells at high concentrations. This data indicates that SP141 disrupts mitosis and cytokinesis, ultimately leading to cell death at higher concentrations.

These results indicate that both Topotecan and SP141 exhibit cytotoxic effects in 92-1 cells, with different potencies and targeting distinct cell cycle phases, supporting the rationale for their synergistic use in UM to enhance their effects at lower doses.

### Topotecan and SP141 synergistically reduce cell viability in 92-1 UM Cells

To evaluate the therapeutic efficacy of the combinatorial effects of topotecan and SP141, we treated 92-1 cells with increasing concentrations of topotecan (2.5 to 25 nM) alongside increasing doses of SP141 (10 to 600 nM) over 72 h. As illustrated in Figure 6A–6E, monotherapy with either topotecan or SP141 resulted in only a limited reduction in cell viability. However, administration of both drugs resulted in a significant dose-dependent decline in cell viability in 92-1 cells. At a 25 nM concentration, topotecan, when paired with 150 nM SP141, reduced cell viability to approximately 25% of control levels and 50-60% of the respective monotherapies, demonstrating significant synergistic effects. A similar response was observed with 5 to 25 nM topotecan in combination with 600 nM SP141, in which the combination reduced cell viability to approximately 25% of control levels and to 25-50% of the corresponding single-agent treatments.

**Figure 6.**
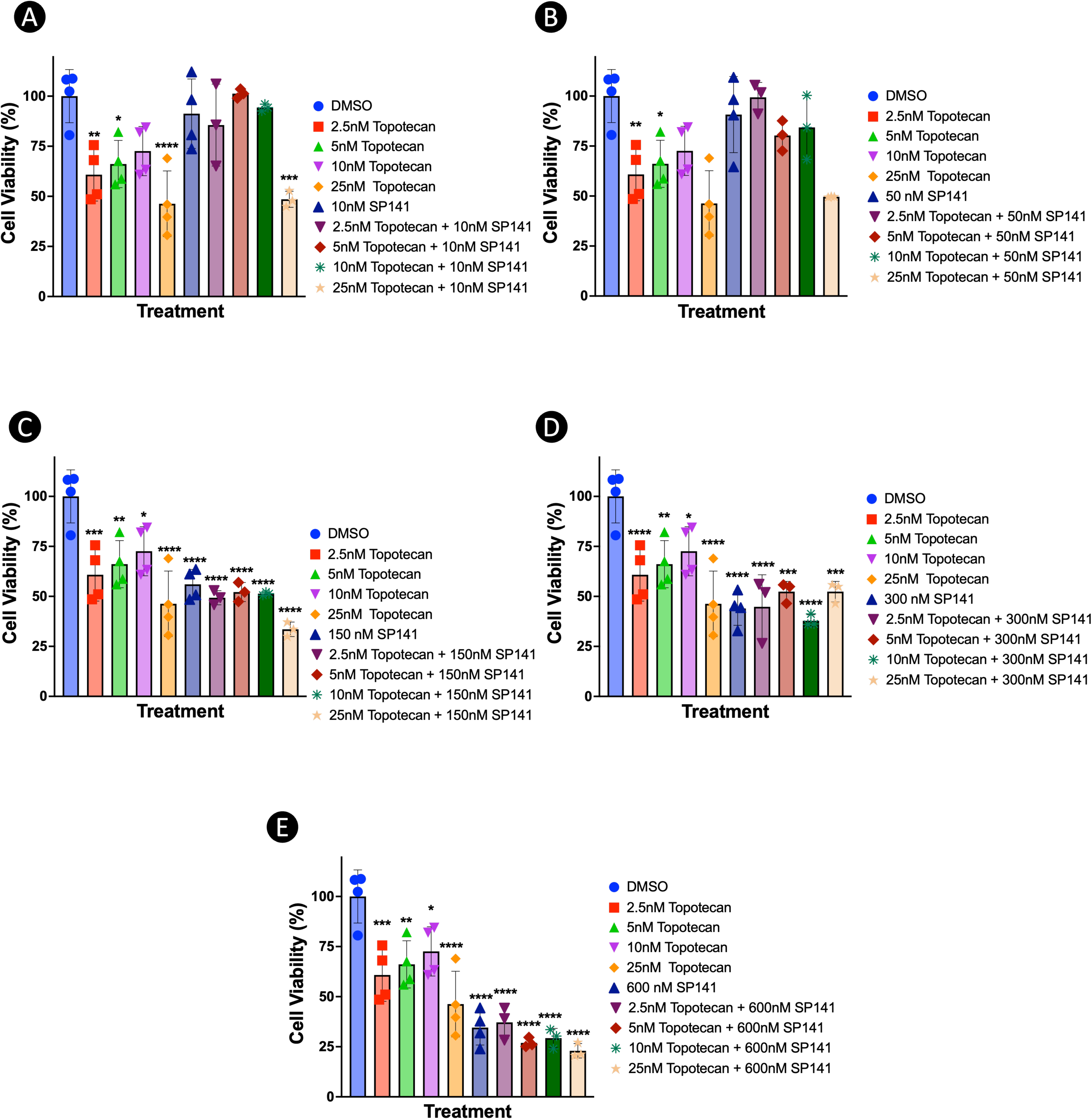
Topotecan and SP141 synergistically reduce cell viability in 92-1 UM cells. Bar graphs (A-E) depicting the percentage cell viability of 92-1 cells, measured by MTT assay after 72-hour treatment with various combinations of Topotecan and SP141. Values are expressed as mean ± standard deviation (SD). Data are represented as mean ± standard deviation (SD). n =3-4 in each group. *, p ≤ 0.05; **, p ≤ 0.01; ***, p<0.001; and ****, p ≤ 0.001 compared with the DMSO group in each fitgure.

The effects of the topotecan and SP141 combination were analyzed using Combenefit software to assess synergistic interactions. Both the Loewe additivity and HSA models were assessed, and heat maps depicting synergism scores were generated (Figures 7A and B). Synergism scores were denoted by blue-shaded cells, indicating a substantial degree of synergism, particularly at the combination of 25 nM topotecan + 150 nM SP141 with Loewe score: +23 and HSA score: +26. Additional synergistic interactions were observed with 25 nM topotecan + 600 nM SP141 (Loewes and HSA score: +16), 5 nM topotecan + 600 nM SP141 (Loewes and HSA score: +12), and 2.5 nM topotecan + 150 nM SP141 (Loewes and HSA score: +10). Notably, lower concentrations of SP141 (10 -50 nM) and topotecan (2.5 -10 nM) in combination showed antagonistic interactions, as indicated by the red/orange shading (Figure 7A and 7 B). This suggests that the topotecan and SP141 combination at lower concentrations sensitizes 92-1 cells to cytotoxic effects, with optimal synergistic effects occurring at 150 to 600 nM SP141 in combination with 2.5 to 25 nM topotecan. This combination strategy suggests sub-IC₅₀ concentrations of both topotencan (independent IC₅₀ 43.8nM) and SP141 (independent IC₅₀ 1324 nM) can synergistically improve anti-cancer efficacy in UM cells.

**Figure 7.**
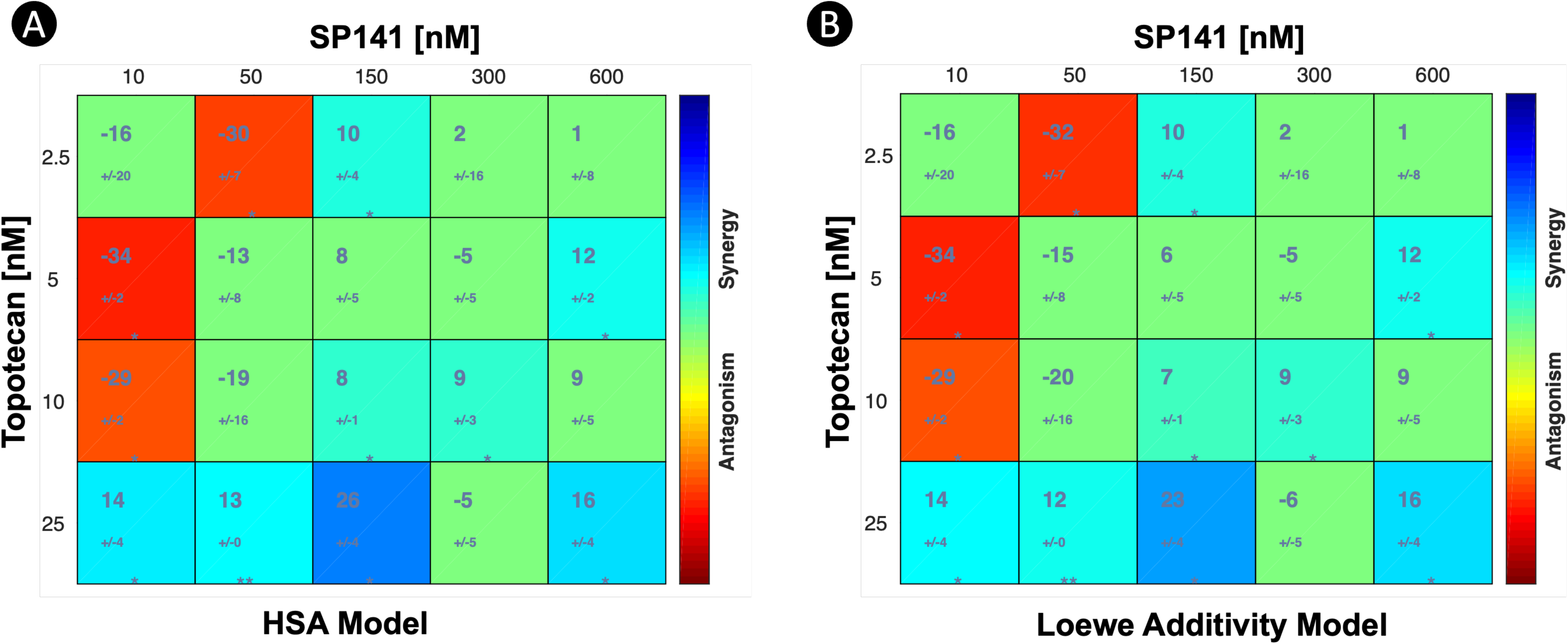
Combenefit analysis suggests synergistic effects of the combined application of topotecan and SP141 in 92-1 UM cells. A) Combination matrix analysis using the HSA (Highest Single Agent) model, indicating regions of synergism and antagonism. B) Combination matrix analysis using the Loewe additivity model, highlighting areas of synergism and antagonism. Data are represented as mean ± standard deviation (SD). n = 3 in each group. *, p ≤ 0.05 and **, p ≤ 0.01.

### Topotecan and SP141 combination causes caspase 3/7 mediated cell death in 92-1 UM cells

To determine whether the combination of topotecan and SP141 enhances cell death, we quantified cell death using Cytotox dyes that monitor membrane integrity in real time and caspase-3/7–dependent apoptosis in 92-1 cells via real-time IncuCyte live-cell imaging. Cells were treated with combination regimens of 25 nM topotecan + 150 nM SP141 and 25 nM topotecan + 600 nM SP141, as well as their respective monotherapies, and were continuously imaged for four days in the presence of Cytotox (Red dye) and Caspase-3/7 (Green dye) reagents. The time course of the Cytotox assay (Figure 8A) shows that monotherapies caused moderate changes in cell death, whereas the combination of topotecan and SP141 produced a greater, earlier increase in cytotoxicity. The 25 nM topotecan + 600 nM SP141 combination produced the most significant effect, with a noticeable difference from monotherapies that began around 40 h and eventually reached a 3-fold increase in cell death by 96 h (Figure 8E). Although 25 nM topotecan alone showed greater cytotoxicity than SP141 monotherapy at early time points (48 to 75 h) (Figure 8D), its effect was comparable to the 25 nM topotecan + 150 nM SP141 combination, which remained significantly higher than both the concentrations of SP141 monotherapies at later stages (Figure 8B–E). These findings indicate that topotecan induces early cytotoxicity, whereas SP141 amplifies and sustains this response in a dose-dependent manner.

**Figure 8.**
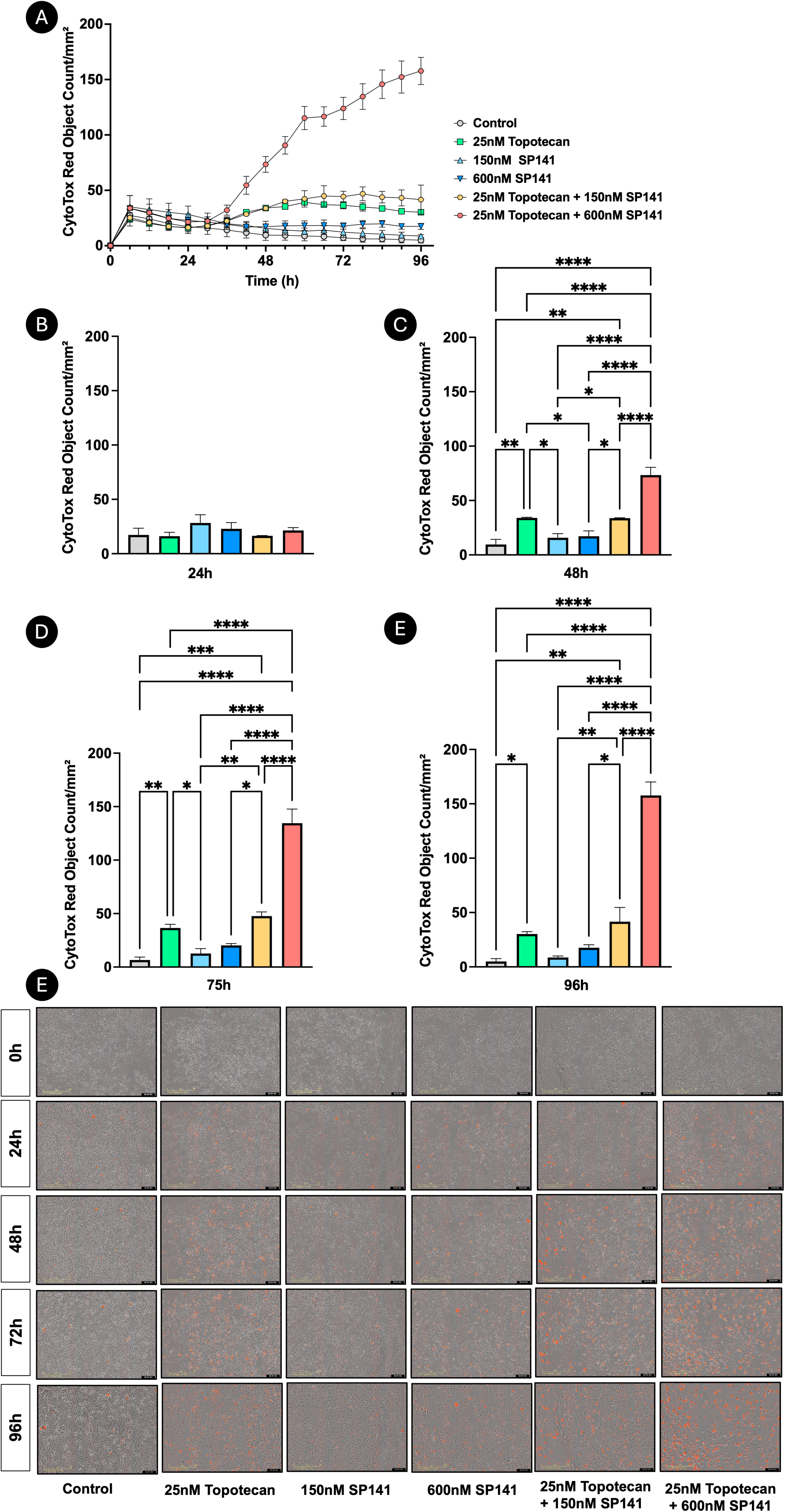
Combined application of topotecan and SP141 induces cytotoxicity in 92-1 UM cells. Time-course analysis of CytoTox Red fluorescence intensity in 92-1 UM cells using Incucyte SX3 live-cell imaging, with cells treated individually with Topotecan, SP141, or combination therapy, demonstrating increased cytotoxicity with combination therapy (A). Bar graphs representing the quantification of Cytotox Red objects at 24 (B), 48 (C), 72 (D), and 96 (E) h. Representative IncuCyte images illustrating CytoTox activity in 92-1 cells following treatment with Topotecan, SP141, or combination therapy. Phase-contrast and red fluorescence images were captured over a 96-hour period to visualize membrane-compromised (dead) cells (E). Images shown are representative fields from each treatment condition at sequential time points. Data are represented as mean ± standard deviation (SD). *, p ≤ 0.05; **, p ≤ 0.01; ***, p ≤ 0.001; and ****,p<0.0001 compared with the control group.

A comparable combinatorial pattern was observed in the Caspase-3/7 apoptosis assay (Figure 9A–9E). The combination of 25 nM topotecan and 600 nM SP141 produced the highest apoptotic activity at all time points, with caspase activation beginning around 40 h and increasing thereafter, consistent with the cell death observed in the Incucyte Cytotox assay. 25 nM topotecan alone induced higher caspase-3/7 activation than SP141 monotherapies at the earlier time point (48 h), whereas SP141 monotherapies showed a delayed increase in caspase activity that became prominent by 96 h (Figure 9E), consistent with a later onset of apoptosis. While caspase activity in SP141 monotherapies continued to increase at later time points, the topotecan monotherapy signal diminished (Figure 9D, E), suggesting differential patterns of apoptotic induction. The 25 nM topotecan + 150 nM SP141 combination showed enhanced apoptotic activity beginning at 48 h compared with monotherapy treatments (Figure 9A, C). Also, the 25 nM topotecan + 600 nM SP141 combination consistently produced the strongest apoptotic response, exceeding both monotherapies and the lower-dose combination at all time points studied.

**Figure 9.**
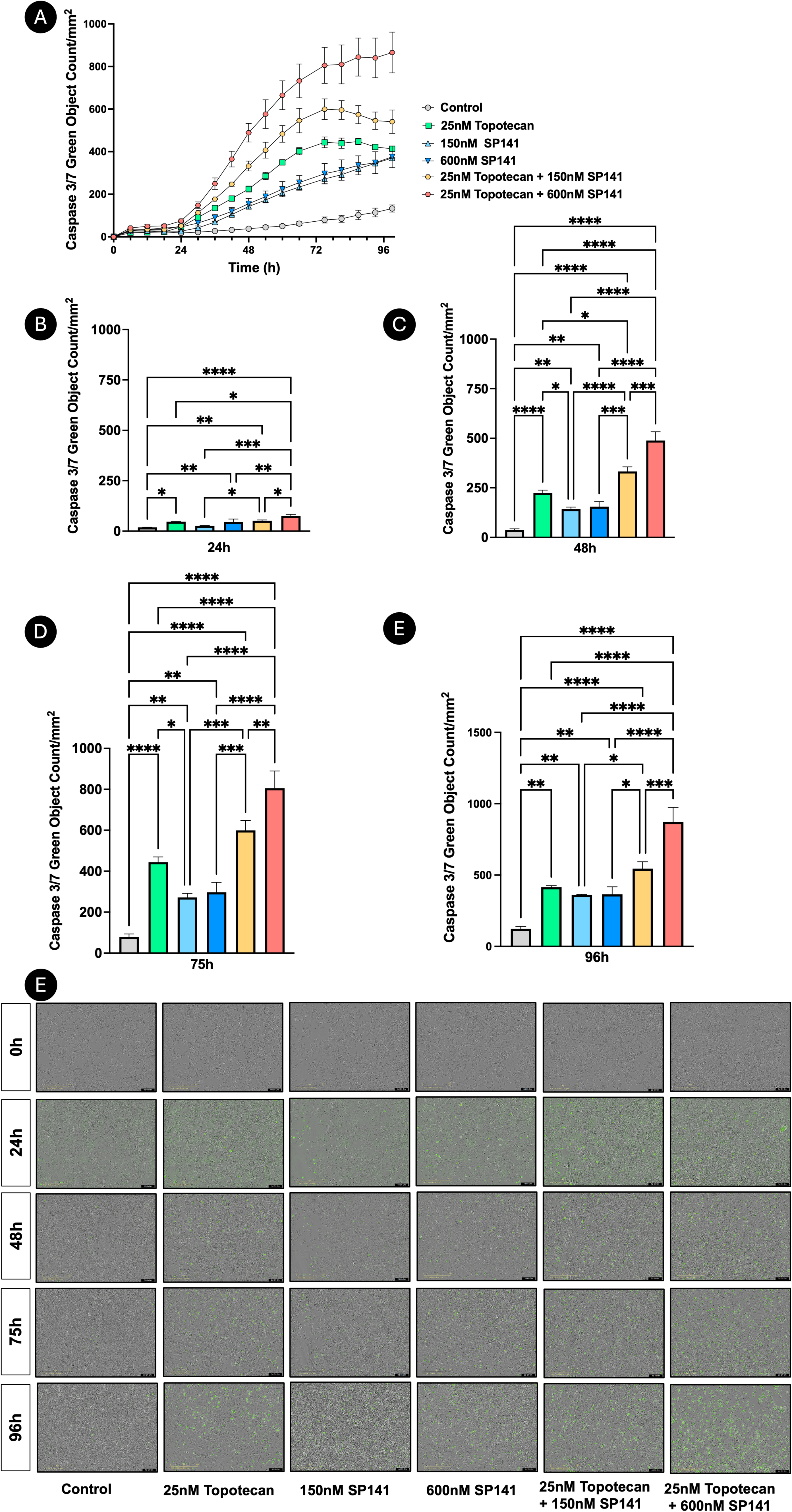
Topotecan and SP141 combination suggests caspase 3/7 mediated cell death in 92-1 UM cells. A) Time-course analysis of Caspase 3/7 Green fluorescence signal in 92-1 UM cells was done using Incucyte SX3 live cell imaging, cells treated with Topotecan, SP141, or combination therapy, demonstrating increased cytotoxicity with combination treatment (A). Bar graphs depicting quantification of Caspase 3/7 green objects at 24 (B), 48 (C), 72 (D), and 96 (E) h. Representative IncuCyte images illustrating cytotoxicity in 92-1 cells following treatment with Topotecan, SP141, or combination therapy. Phase-contrast and Green fluorescence images were captured over a 96-hour period to visualize membrane-compromised (dead) cells. Images shown are representative fields from each treatment condition at sequential time points. Data are represented as mean ± standard deviation (SD). *, p ≤ 0.05; **, p ≤ 0.01; ***, p ≤ 0.001; and ****,p<0.0001 compared with the control group.

Overall, these findings indicate that topotecan triggers apoptosis earlier, whereas SP141 produces a more delayed apoptotic effect. When administered in combination, topotecan and SP141 produce an accelerated, markedly amplified caspase-3/7–mediated apoptotic response, which corresponds to decreased membrane integrity observed in the Cytotox assay. Both early and sustained cell death highlight the potential of synergism between topotecan and SP141 and suggest an effective strategy for promoting apoptosis in 92-1 cells.

### Topotecan and SP-141 exhibit limited toxicity to retinal pigment epithelium

We tested the tolerability of Topotecan and SP141 in ARPE-19 cells, a human retinal pigment epithelial (RPE) cell line that forms the outer blood-retinal barrier (BRB), to assess BRB integrity as the tumor develops beneath it (Fig 10A-D). To assess the effects of topotecan and SP141 on ARPE-19 cells, we treated cells with different concentrations of topotecan (10 to 5000 nM) and SP141 (250 to 20,000 nM) for 72 h. The media was collected and analyzed using MTT and LDH assays to determine cell viability and membrane integrity/cytotoxicity.

**Figure 10.**
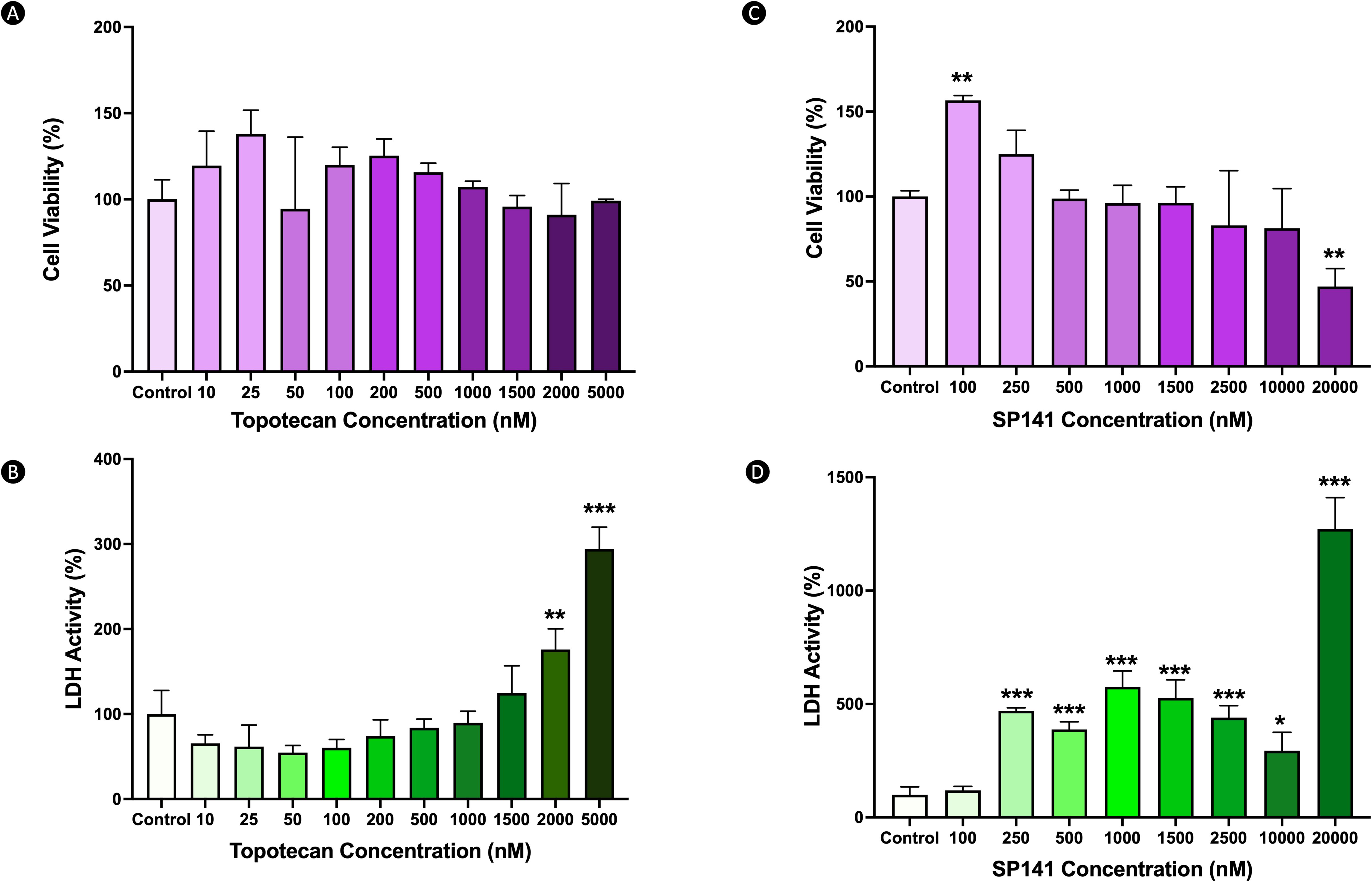
Topotecan and SP-141 demonstrate minimal toxicity to ARPE-19 cells. Bar graphs represent the MTT (A) and LDH (B) assays performed on ARPE-19 cells treated with several concentrations of Topotecan for 72 h. LDH cytotoxicity assay of ARPE-19 cells treated with Topotecan for 72 hours. Similarly, SP141 was treated with ARPE-19 cells at multiple concentrations to assess drug tolerability for 72 hours using MTT (C) and LDH (D) assays. Data are represented as mean ± standard deviation (SD). *, p ≤ 0.05; **, p ≤ 0.01; and ***,p<0.001 compared with the control group.

The MTT assay in ARPE-19 cells after topotecan treatment shows stability across all tested concentrations (Fig 10A). The LDH assay indicates that LDH is retained within viable ARPE-19 cells at all doses except at high concentrations (2000 nM and 5000 nM) after 72 h of treatment (Fig 10B). MTT assay on ARPE-19 cells treated with SP141 shows relative stability across all concentrations with a small but significant increase in metabolic viability observed at 100nM and a decrease in metabolic viability at 20,000 nM SP141 at 72 h (Fig 10C). The LDH assay on media from ARPE-19 cells treated with SP141 shows decreased membrane integrity above 250 nM after 72 h of treatment, with increased cell death at higher concentrations (Fig 10D).

These findings indicate that ARPE-19 cells tolerate low-to-moderate concentrations of topotecan and SP141, with decreased viability at higher doses, suggesting that sub-IC₅₀ levels may be safe for managing UM.

## Discussion

UM represents the most aggressive primary intraocular tumor in adults. Approximately half of the patients develop metastatic disease, predominantly affecting the liver in 80–90% of cases. Once liver metastasis develops, the prognosis is poor, with a median survival of 6 to 12 months and a five-year overall survival rate of less than 5%.^11,37,38^ Currently, tebentafusp-tebn (brand name: Kimmtrak) is the only FDA-approved systemic biphasic T-cell engager therapy for uveal melanoma in HLA-A*02:01–positive patients, whereas melphalan delivered via hepatic perfusion is approved for patients with unresectable hepatic metastases.^39,40^ This highlights the importance of developing new, targeted strategies to address this clinical challenge. The current study examines the therapeutic potential of topotecan, a known topoisomerase-1 inhibitor,^16^ and SP141, an MDM2 inhibitor,^41,42^ in 92-1 UM cells. Reactivation of p53 is considered the most effective way to combat cancer, as it can directly inhibit tumor growth and also activate immune cells, aiding in tumor clearance.^43^ The rationale for combining topotecan and SP141 lies in their complementary mechanisms of action that converge on the p53 pathway. Topotecan induces DNA damage, activating p53 signaling;^44^ however, p53 also transcriptionally upregulates MDM2, creating a negative feedback loop that attenuates p53 activity.^45^ Targeting MDM2 with SP141 may therefore disrupt this feedback regulation, resulting in enhanced and sustained p53 activation^46^ and thereby triggering cell death in the UM cells. Anticancer drug killing potency profiles were assessed to understand the antiproliferative and cytotoxic effects of these drugs at independent doses. Additionally, we subsequently developed a suitable strategy to evaluate the synergistic effects of these drugs on 92-1 cells. For the first time, the study provided sub-IC_50_ values for multi-target cytotoxic effects of Topotecan and SP141 acting synergistically in 92-1 UM cells.

### Potential therapeutic effects of Topotecan and SP141 for UM

Topotecan has been used as an FDA-approved chemotherapy drug ^47–49^ for ovarian cancer and in combination with cisplatin for small-cell lung cancer and cervical cancer. It is in the preclinical investigation phase for colorectal cancers.^50^ In recent years, topotecan has gained popularity in ophthalmology for treating retinoblastoma,^51–53^ either as monotherapy or in combination, demonstrating safety, efficacy, and good ocular tolerability.^54,55^ However, studies evaluating topotecan in UM are limited to a single *in vitro* study demonstrating its efficacy in combination with RITA and Nutelin-3, inducing apoptosis in UM cell lines.^26^ SP141, on the other hand, is among the new MDM2 inhibitors still in preclinical or clinical development. While first-generation MDM2 inhibitors such as Nutlin-3a, RG7112, and RG7388 block p53 degradation by competitively binding to the p53-binding pocket of MDM2, SP141 acts distinctively by promoting MDM2 autoubiquitination and proteasomal degradation. In addition, it reactivates p53 and exerts p53-independent antitumor effects, thereby broadening its therapeutic potential.^25,56–58^ Preclinical studies of SP141 have demonstrated antitumor activity *in vitro* and *in vivo* against breast, pancreatic, and neuroblastoma cancers.^41,42,46^ To our knowledge, the therapeutic use of SP141 has not been explored in UM.

UM typically retains a wild-type *TP53* genotype,^59^ unlike other cancers. However, it exhibits a high propensity for hepatic metastasis. Consequently, strategies aimed at restoring p53 activity, while targeting metastatic progression, are of increasing translational significance. MDM2 is known to play a role in metastasis by regulating epithelial-mesenchymal transition (EMT),^60,61^ and its knockdown has been shown to reduce tumor metastasis.^62^ MDM2 inhibitors such as RG7388 (Idasanutlin) have shown potential to reduce metastatic disease in breast cancer.^63^ Similarly, APG115 (Alrizomadlin) has shown antitumor activity in advanced solid tumors.^64^ Recently, SP141 has been reported to suppress metastatic spread in other cancers,^57^ making it an attractive candidate for further exploration in the context of UM, where metastasis to the liver remains the leading cause of mortality.

Our results demonstrate that combinatorial treatment with topotecan and SP141 induces dose-dependent cytotoxicity in 92-1 UM cells, as evidenced by multiple cellular assays that designate distinct IC_50_ values for these compounds. The IC_50_ values for topotecan and SP141 in 92-1 UM cells after 72 h of application were 43.8 nM and 1.324 µM, respectively. While topotecan has exhibited similar effects in retinoblastoma,^54,55^ evidence supporting its use in ocular tissues, especially in UM, remains limited. To date, only one prior study has reported growth inhibition by topotecan, with an IC₅₀ of 25.78 nM.^26^ This IC₅₀ is slightly lower than the value we observed in our study. However, the variation may be attributable to differences in experimental conditions, assay type, or exposure duration. The IC_50_ values for SP141 have not been reported in ocular cancers. Previous studies on pancreatic and breast cancers reported IC_50_ values ranging from 0.3 μM to 13 μM.^41,42^ Similarly, work on glioblastoma models has suggested an IC_50_ range of 0.03 μM to 3.8 μM.^56^

Our study provides the first evidence of SP141’s potency in UM cells. The observed reduction in cell growth and proliferation can be attributed, at least in part, to disruption of normal cell cycle progression and phase-specific cell cycle arrest. Consistent with previous findings,^65,66^ our study shows that low concentrations of topotecan arrest 92-1 cells in the G2/M phase, while higher doses arrest cells in the S phase. Further, we describe that SP141 arrests the UM cell line in the G2/M phase of the cell cycle. These findings suggest that the cytotoxic effects of both drugs are mediated not only by direct inhibition of proliferation but also by perturbations of checkpoints that compromise cell survival.

### Synergism of Topotecan and SP141 imparts multi-target therapy for UM

The potency profiles and multi-targeted mechanisms of action of topotecan and SP141 offer an opportunity to develop combination therapies at sub-IC_50_ concentrations, demonstrating improved efficacy and enhanced cytotoxicity in 92-1 cells. The data from cellular assays (Figs 6 and 8) and subsequent Combenefit analysis (Loewe additivity and HSA models; Fig 7) indicated maximal synergy between the two drugs at 25 nM topotecan + 150 nM SP141, followed by 25 nM topotecan + 600 nM SP141 combination. In fact, the higher-dose combination of 25 nM topotecan + 600 nM SP141 resulted in increased cell death in the Cytotox assay, as observed in the Incucyte live-cell imaging system. Nevertheless, these effects were observed at concentrations significantly below their respective IC_50_ values: 43.8 nM for topotecan and 1.324 μM for SP141. Additionally, topotecan and SP141 target complementary mechanisms that activate p53, suggesting an additive response in this system. A previous study suggests that replication stress induced by topotecan-mediated DNA double-strand breaks triggers the ATM–CHK2–p53 signaling axis, representing a cellular stress response.^67^ At the same time, SP141 promotes MDM2 degradation, thereby stabilizing and increasing intracellular p53 levels.^41^ Our findings suggest that both drugs interfere with standard checkpoints: topotecan causes dose-dependent cell cycle arrest in the S and G2/M phases, while SP141 induces G2/M arrest. This unique combination enhances checkpoint stalling, thereby enabling multi-target activity. Targeting these mechanisms could improve tumor cell sensitivity and potentially augment apoptotic responses.^26,68^ These results provide a strong rationale for combinatorial therapy, in which drugs with complementary mechanisms and non-overlapping potency profiles have been shown to improve efficacy and reduce toxicity compared with monotherapy.^69^

Topotecan has been reported to induce apoptosis by activating effector caspase-3 and caspase-7 in multiple cancer models,^70,71^ including high-grade gliomas, where caspase-3 activation follows DNA damage–induced replication stress.^70^ Similarly, SP141 has been shown to induce apoptotic cell death by activating effector caspases in pancreatic and breast cancer models.^41,42^ Although the combination of topotecan and SP141 has not been previously explored in ocular tissues, both drugs have demonstrated enhanced apoptotic activity when combined individually with other therapeutic agents. For example, topotecan has been shown to increase caspase-3/7 activity in leukemia cells when combined with amlodipine^72^ , and metronomic administration of topotecan with docetaxel in prostate cancer models resulted in increased caspase-3/7–mediated apoptosis.^73^ In addition, increased apoptotic cell death has been reported in brain tumor models when sub-IC₅₀ doses of SP141 were combined with temozolomide.^56^

Notably, synergistic interactions between DNA-damaging agents and MDM2 inhibitors have been documented in multiple cancers, including UM and neuroblastoma, where nutlin-3 in combination with topotecan significantly enhanced apoptosis compared with monotherapy.^26,74^ These studies reported increased activation of effector caspases, including cleaved caspase-3, supporting a caspase-dependent mechanism of apoptosis. Similar findings have been reported in acute myeloid leukemia, where MDM2 inhibition with Nutlin-3 augmented the effects of Cytarabine,^75^ and in glioblastoma, where Nutlin-3a enhanced the cytotoxicity of Temozolomide.^76^ Consistent with these findings, our study showed that SP141 synergizes with topotecan to suppress cell viability and to accelerate the onset of caspase-3/7–mediated apoptosis in 92-1 cells. Importantly, this enhanced apoptotic response was achieved at sub-IC₅₀ concentrations. This combinatorial strategy shows potential to use lower doses of both drugs, which may help reduce toxicity.

### Tolerability assessment of the Topotecan and SP141 therapy

Furthermore, our study examined the cellular responses of non-cancerous retinal pigment epithelial cells to treatment with topotecan and SP141 to assess potential off-target or cytotoxic effects. In ARPE-19 cells, treatment with either drug depicted limited cytotoxicity across multiple drug concentrations. SP141 induced mild toxicity at higher doses (> 250 nM). No significant cytotoxic effects were observed at lower concentrations, suggesting that a combinatorial drug strategy at sub-IC_50_ levels may be tolerated by nonmalignant retinal pigment epithelial cells. This data supports a previous study demonstrating that topotecan exhibits minimal toxicity at low doses in ARPE19 cells.^77^ This is the first to document the effect of SP141 in ARPE-19 cells. Other MDM2 inhibitors induced minimal apoptosis in ARPE-19 cells, except when administered at substantially higher concentrations.^78^ Our data suggest that topotecan and SP141 are well tolerated by ARPE-19 cells at low doses, indicating no off-target effects.

### Limitations of the study

A limitation of the present study is that all findings were obtained from a single UM cell line (92-1). Nevertheless, 92-1 UM cells were chosen because 85-90% of UMs originate in the choroid and share the genetic, phenotypic, and metastatic characteristics of primary human UMs. Further validation in additional uveal melanoma cell lines derived from the uvea and beyond is recommended. The toxicity assessment in ARPE-19 cells does not fully reflect *in vivo* conditions, but it lays a foundation for future *in vivo* safety studies. However, our findings provide initial evidence of synergism between topotecan and SP141, which may represent a promising multi-targeted therapeutic approach for UM management.

### Summary

In summary, our study provides strong evidence for the combinatorial use of topotecan and SP141 in 92-1 cells as a potential treatment for UM. We provide the first evidence that the MDM2 inhibitor SP141 exhibits antitumor activity in UM cells and that its combination with topotecan yields significant synergistic effects. Both drugs, individually, reduced cell proliferation and viability, with distinctive phase-specific cell cycle arrest, and, in combination, at sub-IC_50_ doses in 92-1 UM cells.

## Acknowledgements

This work was supported by NIH R01 EY029795 to SSC. Part of this work was presented at the Association for Research in Vision and Ophthalmology (ARVO) 2025 meeting.

---

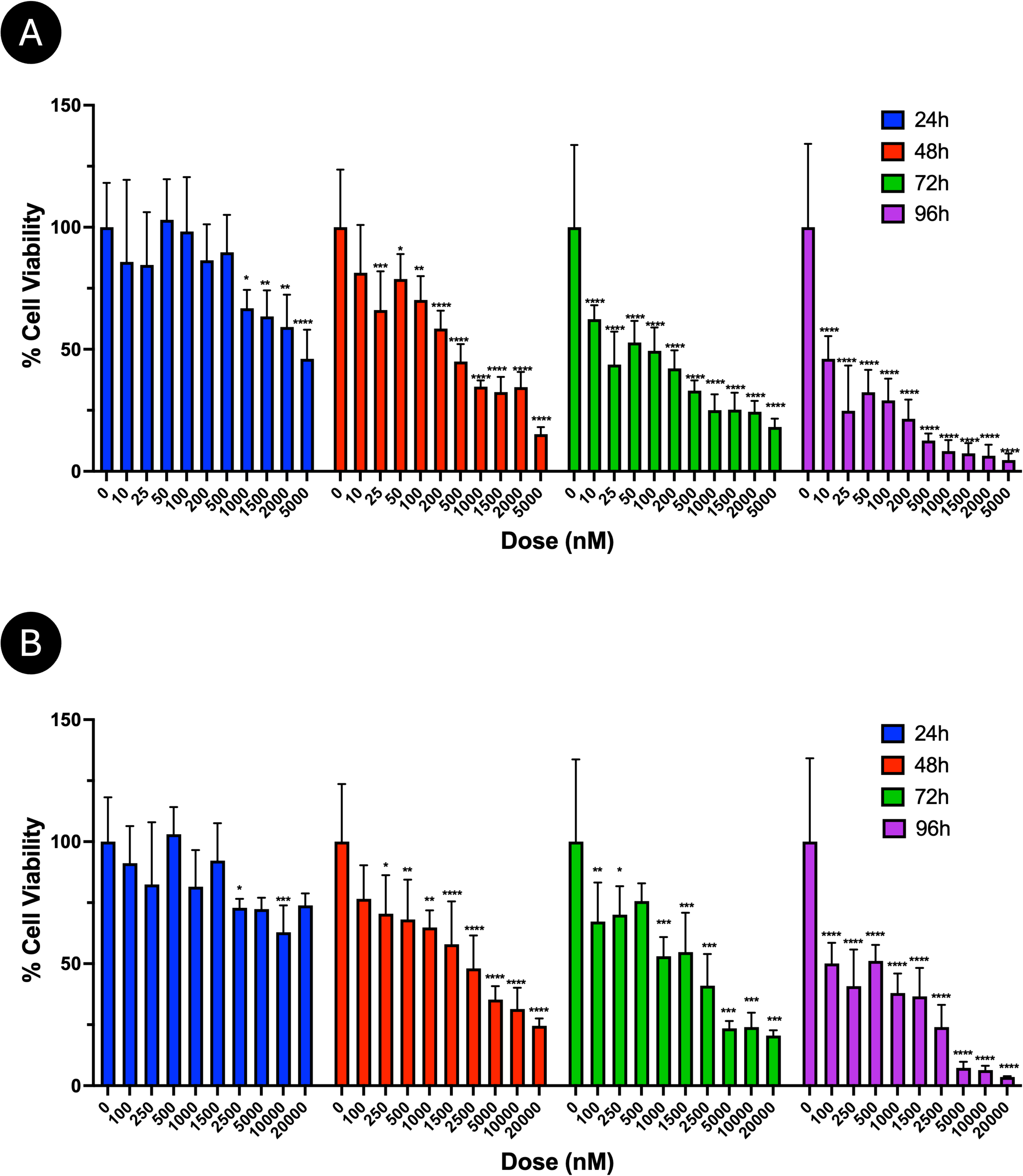

