## Supplementary Figure 1 for "Topotecan Synergizes with SP141 in the Management of Uveal Melanoma"

### **Supplemental Information**

**A**

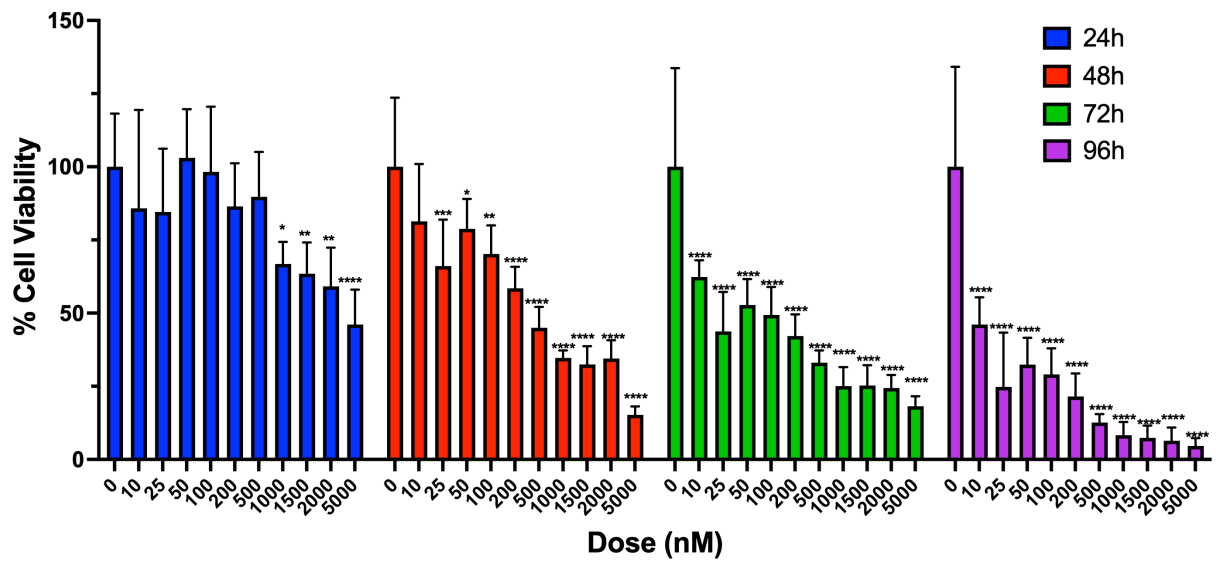

**B**

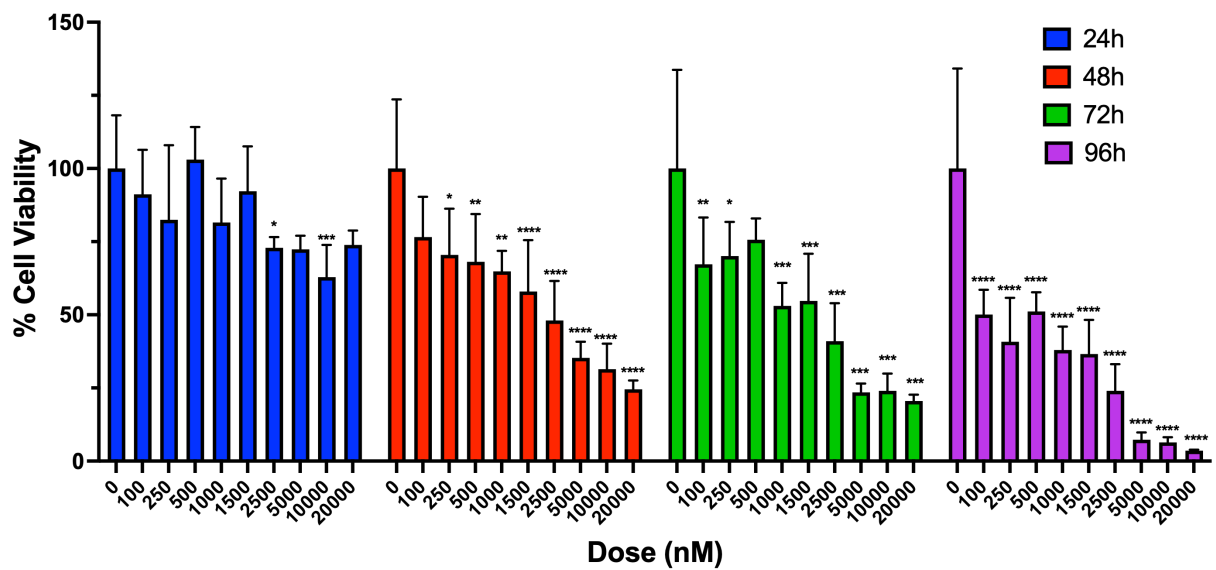

**Supplementary Figure 1. A & B)** Bar graph showing percentage cell viability of 92.1 cells measured by MTT assay after treatment with various concentrations of Topotecan and SP141 for 24, 48, 72, and 96 hours. Values are expressed as mean ± SD ($n = 6$), $P \leq 0.05$ (\*), $P \leq 0.01$ (\*\*), $P \leq 0.001$ (\*\*\*), $P \leq 0.0001$ (\*\*\*\*) indicate statistical significance by Two-way ANOVA.
